# PathwayTheme: identifying and attributing pathway-level axes of variation in omics data

**DOI:** 10.64898/2026.11.22.762686

**Authors:** Aly Abdelkareem, Sofiane Labarre, Sean Nesdoly, Theodore B. Verhey, Shyam V. Menon, Stephen M. Robbins, Donna L. Senger, A. Sorana Morrissy

## Abstract

Exploring gene expression datasets often involves delineating principal axes of variation corresponding to either biological or technical variables. Principal component analysis (PCA) is typically employed as it uses all genes, reflecting global relationships between samples. Existing tools decompose gene expression first and interpret components afterwards. However, gene-level components are sensitive to sparsity and difficult to relate directly to biology, signals carried by low-variance components are easily overlooked, and differential pathway analysis is usually performed separately, on thresholded gene lists. PathwayTheme addresses these limitations by reversing the order of operations: per-sample pathway scores are generated first and then decomposed by PCA, so that each component is defined by pathways and can be tested directly against biological and technical variables. PathwayTheme further incorporates a statistical framework for supervised differential pathway enrichment that does not require thresholded gene lists, with Gene Ontology results summarised into themes. Applied to a published paediatric cancer cohort, PathwayTheme separated technical from biological axes and identified components distinguishing sample types, diagnoses and genetic subgroups, including rare groups represented by low-variance components.

**Availability and implementation:** PathwayTheme is implemented in Python (≥3.10), with an R implementation of the scoring, decomposition and differential-testing steps, and is freely available under the MIT licence at https://github.com/MorrissyLab/PathwayTheme.

## 1 Introduction

Exploratory analyses of gene expression data seek to identify the principal axes of variation, determine whether these correspond to biological or technical variables, and find those that capture the biology of greatest interest, such as molecular subgroups or genetic perturbations [1]. Principal component analysis (PCA) is a common approach because it uses all genes, reflecting global relationships between samples.

Gene-level PCA, however, raises several challenges. First, it is sensitive to sparsity. The fraction of genes above the limit of detection varies between samples with sequencing depth, RNA integrity and library preparation. On a logarithmic scale, the difference between undetected and detected expression is large, so genes detected in some samples but not in others contribute disproportionate variance, and a component can come to reflect detection rate rather than biology [2]. The effect is strongest in single-cell data, but it also arises in bulk cohorts that include different library preparation methods or include degraded samples, and it carries through to any biological interpretation of the affected components. Second, each component is a weighted sum of thousands of genes, so the represented biology must be inferred after decomposition, usually by enrichment analysis of the genes with the largest component loadings. Because genes act together in pathways and many genes participate in several processes, the presence of individual genes among the top loadings is a weak basis for inference, and the result depends on the number of genes selected and on whether both ends of the component are considered. Third, the components of greatest biological interest may explain little of the overall variance. In a cohort that combines tumours, cell lines and normal tissues, or several sequencing protocols, the leading components capture these large differences, whereas subtler distinctions, such as those between tumour subtypes with different genetic drivers, are carried by smaller components that are rarely examined. Tools that relate components to sample variables test each variable as a whole [3–7], which shows whether a component differs among the levels of a variable but not which level it separates. As a result, large cohorts are often analysed through a series of separate, targeted analyses, each designed around a known question (e.g. ref [8]). Exploratory decomposition and differential testing are also usually performed with different tools on different representations of the data, so a component of interest cannot be followed directly by a test on the same features.

Pathway-level representations address the problem of interpretation. Single-sample scoring methods express each sample’s gene expression profile as scores for biological processes [9–14] and, because each score aggregates over many genes, are less sensitive to the detection of individual genes. The resulting score matrix, however, is left to the user to further analyse, typically by clustering or comparisons between predefined groups. Clustering captures only the strongest inter-sample relationships, those carried by the leading principal components, and PCA of pathway scores has been offered mainly as a visualisation [15]. Tools that combine decomposition with pathways instead decompose gene expression first. pcaExplorer and iDEP interpret gene-level components by enrichment analysis of their loadings [7, 16], an approach formalised by PCGSE and GO-PCA [17, 18], and pcaExplorer and several other tools test gene-level components against sample covariates [3–7]. VISION relates gene signature scores to a gene-level latent space in single-cell data [19], and other methods decompose within each gene set [20–22] or project gene sets onto factors derived from the full data [23]. In all of these approaches, the components remain gene-level, so sparsity affects both the components and the pathways inferred from them. Differential pathway analysis is performed separately, usually by over-representation analysis (ORA) of thresholded gene lists, which depends on the threshold and on the balance of upregulated versus downregulated genes between sample groups, and which typically returns many redundant Gene Ontology terms [24–26].

PathwayTheme addresses these limitations by changing the order of the usual operations. Samples are first scored against gene sets, and the resulting pathway score matrix is then decomposed with PCA, so that each component is defined by pathways from the outset and is less sensitive to the detection of individual genes. Because technical factors can also produce coherent pathway-level signals, for example when library preparations capture different classes of transcripts, every component is tested against all recorded biological and technical variables, including the fraction of undetected genes, and against each level of each variable. The same scores support a statistical framework for differential pathway enrichment that does not require thresholded gene lists, with significant terms summarised into Gene Ontology themes. This workflow allows users to identify the component that captures a biology of interest, to determine whether it is confounded with technical factors before defining contrasts, to recover groups carried by low-variance components, and to thematically characterise defined groups without preselected signatures. We illustrate these uses on a published paediatric cohort [8], comparing each with the targeted analysis of the original study.

## 2 Implementation

PathwayTheme is a Python package (≥3.10). Each analysis is defined by a YAML configuration file that reproduces it exactly. The workflow has six steps (Fig. 1A); parameters for all analyses in Section 3 are given in the Supplementary Methods. An R implementation of scoring, decomposition, differential testing and summarisation agrees with the Python implementation to within 2 × 10^−13^ across 38 numerical checks on shared test inputs (Supplementary Methods S13). The association step (Section 2.3) is currently available in Python only.

**Figure 1.**
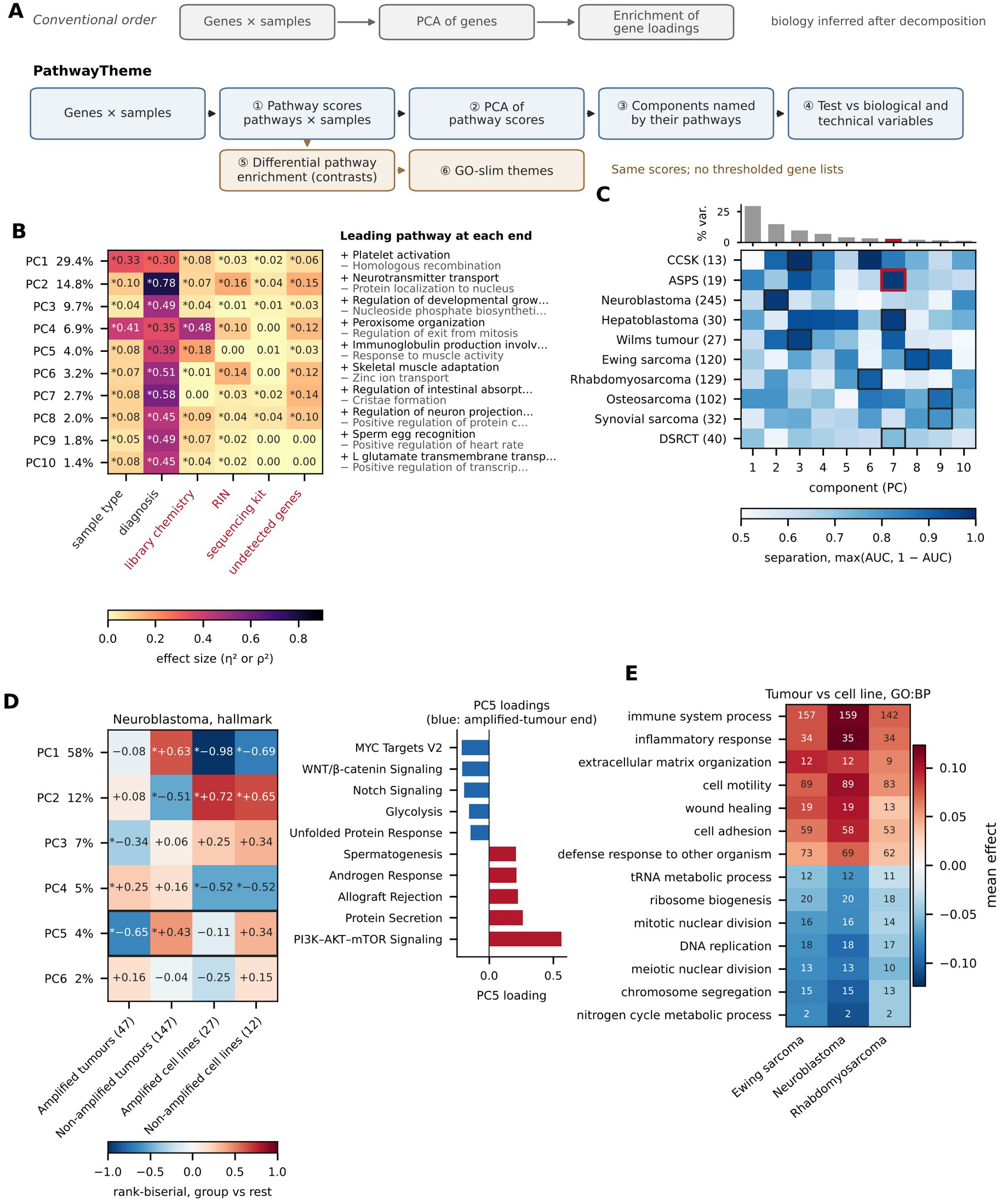
PathwayTheme workflow and use cases in 901 paediatric tumour, normal-tissue and cell-line transcriptomes. (A) PathwayTheme scores samples against gene sets before decomposition (1–4), reversing the conventional order (grey), and tests contrasts on the same scores (5–6). (B) Association of GO:BP components PC1–PC10 with six sample variables, with the leading pathway at each end of each component. Colour, effect size (η^2^ for categorical and ρ^2^ for continuous variables); asterisks, FDR < 0.05 across all 60 tests; red labels, technical variables. ‘Undetected genes’ is the per-sample fraction of protein-coding genes at zero TPM. (C) Separation of each diagnosis with at least ten samples (tumours and cell lines; n in parentheses) from all other tumours and cell lines, as max(AUC, 1 − AUC). Outlined cells mark the best-separating component; the bars show the variance explained by each component. ASPS (red outline) separates on PC7. CCSK, clear cell sarcoma of the kidney; DSRCT, desmoplastic small round cell tumour. (D) *MYCN* amplification in neuroblastoma (194 tumours, 39 cell lines; hallmark collection). Left, rank-biserial correlation of each group against all other neuroblastoma samples on PC1–PC6 (asterisks, FDR < 0.05); PC5 (outlined) separates amplified from non-amplified tumours. Right, the five largest PC5 loadings at each end; blue marks the end occupied by amplified tumours. (E) The 14 GO-slim themes with the largest absolute effect in tumour-versus-cell-line contrasts within Ewing sarcoma, neuroblastoma and rhabdomyosarcoma. Colour, mean effect of the significant member terms of each theme (positive, higher in tumours); numbers, count of those terms. Hallmark results corresponding to panels B and C are in Supplementary Tables S13–S15.

### 2.1 Input and pathway scoring

The input is a gene × sample matrix and a table of sample annotations. Any assay that yields gene-level quantification is accepted, including bulk and single-cell RNA-seq, proteomics, or chromatin accessibility summarised per gene. Each sample is scored against each gene set (step 1), by default with single-sample gene set enrichment analysis (ssGSEA) [12] as implemented in GSEApy [27]. Enrichr ORA [28] and Gene Ontology (GO) slim scoring through GOATOOLS [29] are available as alternatives. Gene sets are filtered by size after restriction to the genes present in the data.

### 2.2 Decomposition and component annotation

Pathway scores are standardised across samples, constant pathways removed, and principal components computed by truncated singular value decomposition [30] (step 2). Sample annotations are not used in the decomposition. Each component is reported with its variance explained, along with pathways with the largest positive and negative loadings (step 3). The number of components annotated is user-defined (default 10; we typically examine up to 20), because low-variance components can separate small sample groups (see Section 3.3).

### 2.3 Association of components with sample variables

Each component is tested against each sample variable (step 4). Categorical variables are tested by the Kruskal–Wallis test, with the rank-based η^2^ = (H − k + 1)/(n − k) as effect size; continuous variables are tested by Spearman correlation, with ρ^2^ as effect size. Levels with fewer than three samples are excluded, and P-values are adjusted across all tests by the Benjamini–Hochberg procedure [31]. Biological and technical variables are tested identically, yielding a single table of associations. The Kruskal–Wallis test indicates whether a component differs among the levels of a variable, but not which level it separates. Each level is therefore also compared with all other samples by the Mann–Whitney test, with the rank-biserial correlation as a signed effect size [32]. Rescaled to [0, 1], this effect size equals the area under the receiver operating characteristic curve (AUC).

### 2.4 Differential testing and theme summarisation

The supervised branch uses the same pathway score matrix (step 5). Contrasts compare one level with all others, with a named reference, or with another level. For each pathway, the effect is the difference in mean score between groups. Significance is assessed by Welch’s t-test, the Mann–Whitney test or a moderated t-statistic with an empirical Bayes variance prior [33, 34], and P-values are adjusted within each contrast by the Benjamini–Hochberg procedure. The moderated t-statistic is a Python port of limma’s variance-prior estimation. Across 108,195 tests from 24 contrasts, it matched limma::eBayes to within 1.3 × 10^−11^ and altered no significance call (Supplementary Fig. S1).

Differential analysis of GO biological process (GO:BP) terms typically yields thousands of overlapping significant terms. Theme summarisation (step 6) maps each term to its most specific ancestor in the generic GO slim, following *is_a*, *part_of* and *regulates* relations [35]. Each theme is reported with the mean effect of its significant terms and the numbers of terms increased and decreased. Non-significant and unmapped terms are reported rather than discarded.

### 2.5 Scope and limitations

Only bulk RNA-seq is evaluated here; other gene-level measurements, and decomposition within user-defined sample partitions, are supported but untested. Differential tests do not adjust for covariates; where the association analysis indicates confounding, contrasts can be stratified, or tested with covariate-adjusted models in other tools. Because GO:BP terms overlap, the number of significant terms overstates the number of independent findings; theme summarisation reduces this redundancy in presentation, not in inference.

## 3 Use cases

### 3.1 Cohort and reference analysis

We applied PathwayTheme to the 901 publicly available transcriptomes of the paediatric solid-tumour cohort of Brohl et al. [8], comprising 654 tumours, 118 normal tissues and 129 cell lines (Supplementary Table S1). The cohort combines several studies, spans rare and abundant diagnoses (5 to 245 samples per diagnosis), includes tumours and disease-matched cell-line models, and normal tissues from multiple organs. Library chemistry is partly confounded with sample type and diagnosis, so technical and biological signals are intermixed.

The original analysis addressed four questions, each with a separate analysis designed in advance [8]. Confounding by library chemistry was handled by restricting comparisons to samples of matching preparation; differences in immune content between diagnoses were assessed with 24 predefined immune signatures; the resemblance of cell lines to tumours was examined one candidate therapeutic target at a time; and the relationship between *MYCN* amplification and the immune microenvironment was examined with selected immune signatures and checkpoint genes. The following sections address the same questions within a single workflow on GO:BP scores, without prior specification of signatures, comparisons or confounders.

### 3.2 Separating biological from technical axes

Rather than restricting comparisons to samples of matching preparation, PathwayTheme reports the association of every component with every recorded variable (Fig. 1B; Supplementary Table S2), showing how biological and technical variables align with major and minor axes of variation. The fraction of undetected genes was at most weakly associated with any component (ρ^2^ ≤ 0.15), indicating that differences in detection rate between samples did not dominate any component of the pathway decomposition.

The leading component (PC1) was associated with sample type and diagnosis rather than library chemistry, ordering samples from cell lines through tumours to normal tissues. Platelet activation, cell adhesion and chemotaxis had the largest loadings at the end occupied by normal tissues, and DNA repair and homologous recombination at the end occupied by cell lines, identifying PC1 as an axis of non-malignant tissue content (Supplementary Fig. S2, Table S3).

Library chemistry was associated mainly with the fourth component (PC4), which was also associated with sample type, because all cell lines and most normal tissue libraries were ribosomal RNA-depleted whereas most tumour libraries were poly(A)-selected. To determine whether PC4 reflects sample type beyond chemistry, the sample-type test was repeated within each chemistry. Among ribosomal RNA-depleted samples alone, sample type remained strongly associated with PC4 (Supplementary Table S4), so this component carries sample-type differences that chemistry does not completely explain. In contrast, all Wilms tumours and clear cell sarcomas of the kidney (CCSK) were sequenced after ribosomal RNA depletion, and all hepatoblastomas after poly(A) selection, precluding the separation of technical from biological effects for these diagnoses (Supplementary Fig. S3, Table S5). This overview shows, before any contrast is defined, which comparisons are directly interpretable and which require stratification.

### 3.3 Locating small diagnostic groups

The leading components of a heterogeneous cohort typically represent its largest transcriptional differences, here sample type (PC1) and library chemistry (PC4). In contrast, tumour type (diagnosis) was associated with every component (Fig. 1B). PathwayTheme therefore also tests each individual diagnosis against every component and reports the component best separating it from others (Fig. 1C; Supplementary Tables S6 and S7). Seven of the ten more common diagnoses (≥10 samples each) were strongly separated by a single component, indicating that each has a distinct pathway-level profile. Alveolar soft part sarcoma (ASPS), for instance, separated on PC7. This component explains less than 3% of the cohort variance and lies beyond a conventional scree-plot cut-off, so it would not be examined in a typical analysis. Further exploration of this component in related work showed that its pathways, including mitochondrial, autophagy and muscle-related processes, are strongly concordant with established ASPS biology [36], showing that low-variance components can capture disease-specific biology.

### 3.4 Assessing the fidelity of cell-line models

Cell lines are central experimental tools for investigating and perturbing tumour biology. Grown *in vitro*, they lack the tumour microenvironment but should retain tumour-cell-intrinsic programmes. PathwayTheme assesses the fidelity of cell lines to their tumours of origin at the pathway level with unsupervised and supervised approaches, supporting the validation of new and existing models [36].

In support of this, the unsupervised pathway-centric sample stratification demonstrated clean delineation of tumour-intrinsic activities in ASPS tumours and cell lines (PC7) from those arising *in vivo* from the microenvironmental compartment (PC1; Section 3.2), indicating PathwayTheme can resolve tumour cell-intrinsic biology even when the overall signals are low within a cohort (<3% of total variance), similar to findings in ref [36].

Supervised analyses enable further investigation of tumour versus cell line differences by summarising distinct vs common pathways thematically. For instance, in Ewing sarcoma, neuroblastoma and rhabdomyosarcoma, tumours showed higher immune, extracellular matrix and adhesion themes, and cell lines higher DNA replication and chromosome segregation themes (Fig. 1E; Supplementary Table S8). These contrasts are confounded with library chemistry, although sample-type differences persist within ribosomal RNA-depleted samples (Section 3.2).

### 3.5 Locating the axis of a known biological distinction

The transcriptional effect of genetic drivers is frequently of central interest in tumour biology. Determining whether, and on which component, a genetic event produces a measurable transcriptional shift is a first step in assessing the nature and magnitude of its impact.

Using both unsupervised and supervised approaches, we investigated the impact of *MYCN* amplification in neuroblastoma samples including 194 tumours (47 *MYCN*-amplified and 147 non-amplified) and 39 cell lines (27 amplified, 12 non-amplified). In the unsupervised analysis, *MYCN* amplification stratified tumours on the fifth component (PC5; 4% of variance), orthogonal to separation of cell lines from tumours (PC1), indicating agreement in tumour cell-intrinsic transcriptional shifts (Fig. 1D). PC5 loadings place MYC targets (hallmark MYC targets V2 [37]), WNT/β-catenin and Notch signalling as high in amplified samples, opposite of PI3K–AKT–mTOR signalling, protein secretion and allograft rejection in non-amplified samples.

Supervised analyses were consistent, identifying immune themes as the most decreased in amplified tumours (Supplementary Fig. S4, Tables S9–S11), in agreement with the immune differences that the original study validated at the protein level [8]. The same immune themes were decreased in *MYCN*-amplified cell lines, which lack immune infiltrate, indicating a tumour-cell-intrinsic contribution.

### 3.6 Differential enrichment of pathways among sample groups

Pathway-level differences are commonly tested by ORA of differentially expressed gene lists, for example with g:Profiler and g:SCS correction [38]. Typically, ORA requires thresholds on significance and fold change, discards the magnitude of changes below them, and tests increased and decreased genes as separate lists. Consequently, results depend on the thresholds, on the size of each list and on the balance between them [24–26]. We compared ORA with PathwayTheme for the three pairwise contrasts among tumours, cell lines and normal tissues (Supplementary Fig. S5, Table S12). Every theme identified by ORA was also identified by PathwayTheme, whereas ORA identified fewer than half of the themes. The difference arose largely from imbalanced gene lists: in cell lines relative to tumours, 1,614 genes were decreased with FDR < 0.05 and |log_2_ fold change| > 2, but only 54 were increased – too few to detect enrichment among increased genes. Testing per-sample pathway scores removes the dependence on thresholds and list size, enabling detection of modest but coordinated changes across many genes that fall below a gene-level threshold.

## 4 Conclusions

PathwayTheme applies PCA to pathway scores rather than to genes, so that each component is defined by pathways from the outset, and tests every component against all recorded sample variables. In contrast to workflows that decompose gene expression and interpret components afterwards, or that cluster samples on pathway scores, it identifies which axis of variation carries a biology of interest, including axes that explain little variance, and whether that axis is confounded with other biological or technical factors. The same scores support differential pathway enrichment without thresholded gene lists, summarised into Gene Ontology themes. PathwayTheme thereby consolidates analyses that are usually performed separately into a single exploratory workflow, applicable to expression cohorts that combine tumour types, model systems or sequencing protocols. Planned extensions include covariate adjustment in differential testing and evaluation on single-cell and proteomic data.

## Supporting information

Supplementary Data (configuration file)

## Acknowledgements

Use of AI. Large language models were used as a writing and engineering aid, under author supervision throughout. They assisted with drafting and editing the manuscript text, with implementing and refactoring parts of the software, and with building the figures and the reproducibility checks. They were not used to generate, select or interpret data. The authors designed the study, verified the analysis, and take full responsibility for the content of the manuscript.

## Author contributions

Aly Abdelkareem: Conceptualization; Software; Formal analysis; Visualization; Writing – original draft.

Sofiane Labarre: Conceptualization; Software; Formal analysis; Visualization.

Sean Nesdoly: Investigation; Data curation.

Theodore B. Verhey: Investigation; Data curation.

Shyam V. Menon: Investigation; Data curation.

Stephen M. Robbins: Conceptualization.

Donna L. Senger: Conceptualization.

A. Sorana Morrissy: Conceptualization; Supervision; Writing – original draft; Writing – review and editing.

## Data availability

This study analysed the published data of Brohl et al. [8], available from dbGaP (phs000466, phs000467, phs000468, phs000720, phs000768, phs001052 and phs001928) and GEO (GSE89413 and GSE84629). Sample annotation was taken from Supplementary Table S1 of Brohl et al. [8]. The configuration files for every analysis and the supplementary tables are provided as Supplementary Data. The package and the scripts that generate the figures are available at the repository given under Availability and implementation; the scripts for the comparison with ORA (Section 3.6) will be added to the repository before journal submission.

## Supplementary Material

### Supplementary Methods

#### S1 Cohort

The analysis used the 901 publicly available transcriptomes of the paediatric solid-tumour, normal-tissue and cell-line cohort of Brohl et al. [8]: 654 tumours, 129 cell lines and 118 normal tissues (Table S1). We note available data differs from the 935 transcriptomes reported in the original study.

Diagnoses were taken from the sequencing run sheet as OncoTree codes. In the run sheet, three clear cell sarcomas of the kidney are coded CCS and two Ewing sarcomas ELS; Brohl et al. record them as clear cell sarcoma of the kidney and Ewing sarcoma (Table S1). The run-sheet codes were retained, so these five samples form separate levels. They fall below the size thresholds used for interpretation and appear only in the complete results (Supplementary Fig. S1; Tables S16 and S17). Normal tissues and four teratomas carry no diagnosis code.

The *MYCN* status of neuroblastoma samples was read from the diagnosis code of Brohl et al. (Supplementary Table S1A of that study): 74 amplified (47 tumours, 27 cell lines), 159 non-amplified (147 tumours, 12 cell lines) and 12 tumours of unknown status, which were excluded from the *MYCN* analyses.

#### S2 Expression matrix

Gene-level TPM values for 33,967 Ensembl genes were mapped to HGNC symbols with GENCODE v36 [S1], restricted to the 18,573 protein-coding genes and transformed to log_2_(TPM + 1). No batch correction or imputation was applied, because the analysis assesses the variation associated with technical variables.

For each sample, the fraction of protein-coding genes at zero TPM was recorded as a technical variable (‘undetected genes’ in Fig. 1B). Samples with a fraction above 0.50 would have been excluded; the maximum observed was 0.307, so no sample was excluded.

#### S3 Sample variables

Sample type, diagnosis, library type, RIN, sequencing kit and flowcell were taken from the run sheet, in which a blank sample type denotes normal tissue. The run sheet has no entries for normal tissues and is incomplete for some tumours and cell lines. Three fields were therefore completed from the same measurements recorded elsewhere, and Table S1 gives the source of every value. Library type was completed from Supplementary Table S1A of Brohl et al. (from 783 to 901 samples), and RIN from their Table S1B (from 355 to 631 samples; the two sources agree for 322 of the 334 samples recorded in both). Sequencing kit was completed from other samples on the same flowcell, each flowcell carrying a single kit (from 494 to 596 samples). The flowcell of each normal tissue was read from its sample identifier.

The association analysis (Section S6) used diagnosis, sample type, library type, RIN, sequencing kit and the fraction of undetected genes. Flowcell was excluded because most of its 92 levels contain fewer than three samples. The single sample sequenced with the NextSeq500 (2 × 80) v2 kit also fell below this threshold, so sequencing kit was tested on 595 samples in three levels. Flowcell and 15 alignment and library-quality metrics from Table S1B of Brohl et al. were used in the confounding analysis (Section S8).

#### S4 Gene sets and pathway scoring

Two collections were scored: the 50 hallmark gene sets of the Molecular Signatures Database [S2] and Gene Ontology biological process terms [S3]. Gene sets were restricted to the genes present in the matrix and filtered to 10–500 genes, leaving 50 hallmark sets and 5,145 GO:BP terms. Each sample was scored by ssGSEA [12] in GSEApy 1.3.0 [27], with rank normalisation and weight 0.25. Because ssGSEA scores each sample independently, each collection was scored once and all analyses of that collection used the same scores.

#### S5 Decomposition

In each analysis, scores were standardised per pathway across the analysed samples to mean zero and unit population standard deviation; missing scores were set to zero, and pathways with standard deviation ≤ 10^−12^ were removed. Truncated singular value decomposition was computed with scikit-learn [30] (randomised solver; random seed 42), retaining K = min(n − 1, 10) components for n samples. Component scores are the left singular vectors scaled by the singular values, and loadings are the right singular vectors, of unit length. The variance explained by component k is S_k_^2^/(n − 1) for singular value S_k_, reported as a fraction of the total variance of the standardised matrix.

Sample variables do not enter the decomposition. The sign of a component is arbitrary but reproducible for a given input, so results are interpreted by the magnitude of separation and by which end of the component a group occupies. Sections 3.2–3.4 and 3.6 of the main text use the decomposition of all 901 samples in each collection; Section 3.5 uses a separate decomposition of the neuroblastoma samples (Section S11).

#### S6 Association of components with sample variables

Each of the ten components was tested against each of the six variables of Section S3: categorical variables by the Kruskal–Wallis test, with the rank-based effect size η^2^_H_ = (H − k + 1)/(n − k) for statistic H, k levels and n samples, and continuous variables by Spearman correlation, with ρ^2^ as effect size. Levels with fewer than three samples were removed before testing. P-values were adjusted across all 60 tests by the Benjamini–Hochberg procedure [31] (Table S2; hallmark, Table S13).

Each level with at least three samples was also compared with all other samples in which the variable was recorded, by the Mann–Whitney test, with the rank-biserial correlation r as signed effect size [32]. The AUC equals (1 + r)/2. Separation of a diagnosis by a component was summarised as max(AUC, 1 − AUC) and is reported for the ten diagnoses with at least ten samples, among the 777 tumours and cell lines belonging to diagnoses with at least three samples. The best-separating component is the one with the largest |r| (Tables S6, S7 and S15). Separation ≥ 0.90 is described as almost complete; this threshold is descriptive.

#### S7 Chemistry-stratified association

To determine whether a component associated with both sample type and library chemistry retains sample-type structure independent of chemistry, sample type was tested against PC1 and PC4 within each chemistry stratum, in both collections (Table S4). The four recorded chemistries were collapsed to two strata, poly(A) selection (poly(A) and stranded poly(A); 600 samples) and ribosomal RNA depletion (Ribo-Zero and RiboMinus; 301 samples), because the finer distinction leaves groups below the size threshold. This test reports the uncorrected η^2^, the between-group sum of squares divided by the total sum of squares of the component scores, with P-values from the Kruskal–Wallis test and Benjamini–Hochberg adjustment across the 20 tests. Because this estimator differs from the rank-based η^2^_H_ of Section S6, unconditional values in Table S4 differ slightly from those in Tables S2 and S13.

#### S8 Level-wise confounding

Each level of sample type and diagnosis with at least eight samples (13 levels) was compared with each of 20 technical variables: library type, sequencing kit, flowcell, RIN, the fraction of undetected genes and 15 alignment and library-quality metrics (Supplementary Fig. S3; Table S5). Each level was compared with the samples in which its own variable is recorded, so that incomplete annotation does not appear as confounding. For categorical technical variables, the share of the level in its largest stratum was compared with the same share in the subset; a level was flagged when at least 90% of its samples fell in one stratum while the subset baseline was below 90%. For continuous technical variables, the level was compared with the rest of the subset by Cohen’s d [S4], and |d| ≥ 0.8 was flagged.

#### S9 Differential testing

Contrasts compare one level with all others, every level with a named reference, or specified pairs of levels (Table S17). For each pathway, the effect is the difference in mean score, case minus reference, and P-values are adjusted within each contrast by the Benjamini–Hochberg procedure. All analyses reported here used the moderated t-statistic [33, 34]. For pathway i, the pooled residual variance s_i_^2^ on d_i_ = n_1_ + n_2_ − 2 degrees of freedom is shrunk towards a prior variance s_0_^2^ with d_0_ prior degrees of freedom, estimated by fitting a scaled F-distribution to the residual variances by the method of moments, as in limma. The moderated statistic is the effect divided by its standard error computed from the posterior variance (d_0_s_0_^2^ + d_i_s_i_^2^)/(d_0_ + d_i_), referred to a t-distribution on d_0_ + d_i_ degrees of freedom.

The implementation was compared with limma::eBayes [34] by re-running four analyses (both collections; one-versus-rest and pairwise contrasts) through limma on the same scores, samples and design. Over 108,195 pathway tests from 24 contrasts, effects agreed to within 1.7 × 10^−14^, moderated t-statistics to within 1.3 × 10^−11^ and P-values to within 3.5 × 10^−12^; the prior parameters agreed to a relative difference of 9 × 10^−14^, and no pathway changed significance (Supplementary Fig. S1; Table S16).

#### S10 Theme summarisation

GO:BP terms were mapped to the generic GO slim [35]. Each term was assigned to its most specific ancestor in the slim, following *is_a*, *part_of* and the three *regulates* relations; terms in the slim map to themselves, and the biological process root is not used. This yields 68 themes and maps 5,012 of the 7,583 GO:BP terms (66.1%) before size filtering. For each contrast and theme, member terms were divided into significant (FDR < 0.05) and non-significant, and each group is reported with its mean effect and the numbers of terms increased and decreased (Table S8). Unmapped terms are counted and reported. Hallmark results are not summarised.

#### S11 MYCN analysis in neuroblastoma

The 194 neuroblastoma tumours and 39 cell lines of known *MYCN* status were decomposed separately in the hallmark collection, and each of the four groups defined by *MYCN* status and sample type was compared with all other neuroblastoma samples on each component by the Mann–Whitney test (Fig. 1D). Differential testing compared amplified with non-amplified samples within tumours (47 versus 147) in both collections, and within cell lines (27 versus 12) in GO:BP (Supplementary Fig. S4A, B; Tables S9 and S10). A further analysis of all four groups compared amplified with non-amplified samples within each sample type, and cell lines with tumours within each *MYCN* status (Supplementary Fig. S4C; Table S11).

#### S12 Comparison with over-representation analysis

For each of the three pairwise sample-type contrasts (tumour versus normal tissue, cell line versus normal tissue and cell line versus tumour), genes were tested by the moderated t-statistic on log_2_(TPM + 1) with the same samples. Genes with FDR < 0.05 and an absolute log_2_ fold change greater than 2 were divided by direction, and each list was submitted to g:GOSt with g:SCS correction (g:Profiler [38]) against GO:BP (effective domain size, 18,442 genes). The resulting lists contained 986 increased and 17 decreased genes (tumour versus normal tissue), 310 and 605 (cell line versus normal tissue), and 54 and 1,614 (cell line versus tumour). Significant terms were mapped to the 68 themes of Section S10, and a theme was counted as reached if at least one member term was significant (Supplementary Fig. S5; Table S12). Of the 204 theme–contrast pairs, three had opposite signs in the two analyses; in each, the PathwayTheme mean effect was within 0.012 of zero.

#### S13 Software and configuration files

PathwayTheme 0.1.0 requires Python ≥ 3.10 and uses GSEApy, scikit-learn, SciPy and GOATOOLS [29]. Each analysis is specified by a YAML configuration file; the ten files used here are provided as Supplementary Data, with input paths relative to the analysis directory of the repository. hallmark_sampletype.yaml and gobp_sampletype.yaml define the decompositions of all 901 samples, the associations (Tables S2, S3, S7, S13–S15) and the one-versus-rest sample-type contrasts. gobp_tumor_normal.yaml defines the pairwise sample-type contrasts, gobp_matched.yaml the tumour-versus-cell-line contrasts within Ewing sarcoma, neuroblastoma and rhabdomyosarcoma (Fig. 1E), and gobp_diagnosis.yaml the one-versus-rest diagnosis contrasts within tumours. hallmark_nbl_mycn.yaml, gobp_nbl_mycn.yaml and gobp_nbl_mycn_celllines.yaml define the *MYCN* contrasts, and hallmark_nbl_mycn_4way.yaml and gobp_nbl_mycn_4way.yaml the four-group analyses (Fig. 1D; Supplementary Fig. S4).

An R implementation covers scoring, decomposition, differential testing and theme summarisation; the association step is available in Python only. The two implementations are compared by 38 numerical checks on shared synthetic inputs (scripts r/parity/run_python.py and r/parity/compare.R in the repository). The checks cover ssGSEA scores (including inputs with many tied values), over-representation and GO-slim scoring, gene-set coverage, PCA variance, scores, loadings and signatures, all three differential tests with all three contrast forms, the limma variance model, Benjamini–Hochberg correction and theme summarisation. All agree to within 2 × 10^−13^, the largest difference arising in the PCA scores.

## Supplementary Figures

**Supplementary Fig. S1.**
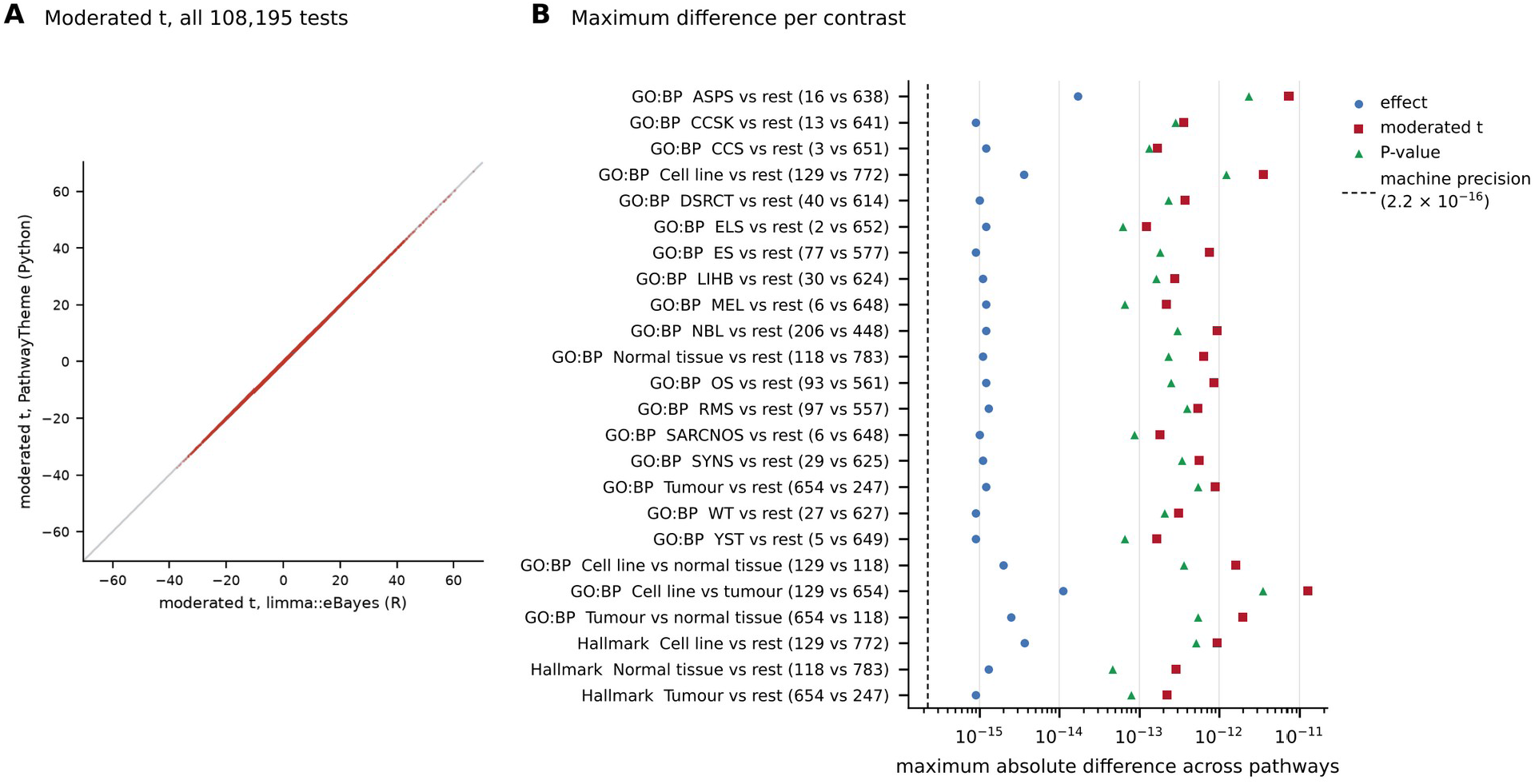
Agreement of the moderated t-test in PathwayTheme with limma::eBayes. (A) Moderated t-statistics from PathwayTheme against those from limma for all 108,195 pathway tests from 24 contrasts. (B) For each contrast, the largest absolute difference between the two implementations across pathways in effect, moderated t-statistic and P-value. The dashed line marks machine precision for double-precision floating-point arithmetic (2.2 × 10^−16^), the smallest relative difference between two numbers that this format can represent; differences of this order are rounding error. Labels give the collection, the contrast and the group sizes (case vs reference). CCS and ELS are run-sheet codes for three clear cell sarcomas of the kidney and two Ewing sarcomas (Supplementary Methods S1). Diagnosis codes: ASPS, alveolar soft part sarcoma; CCS, run-sheet code for clear cell sarcoma of the kidney; CCSK, clear cell sarcoma of the kidney; DSRCT, desmoplastic small round cell tumour; ELS, run-sheet code for Ewing sarcoma; ES, Ewing sarcoma; LIHB, hepatoblastoma; MEL, melanoma; NBL, neuroblastoma; OS, osteosarcoma; RMS, rhabdomyosarcoma; SARCNOS, sarcoma, not otherwise specified; SYNS, synovial sarcoma; WT, Wilms tumour; YST, yolk sac tumour. Values are in Table S16.

**Supplementary Fig. S2.**
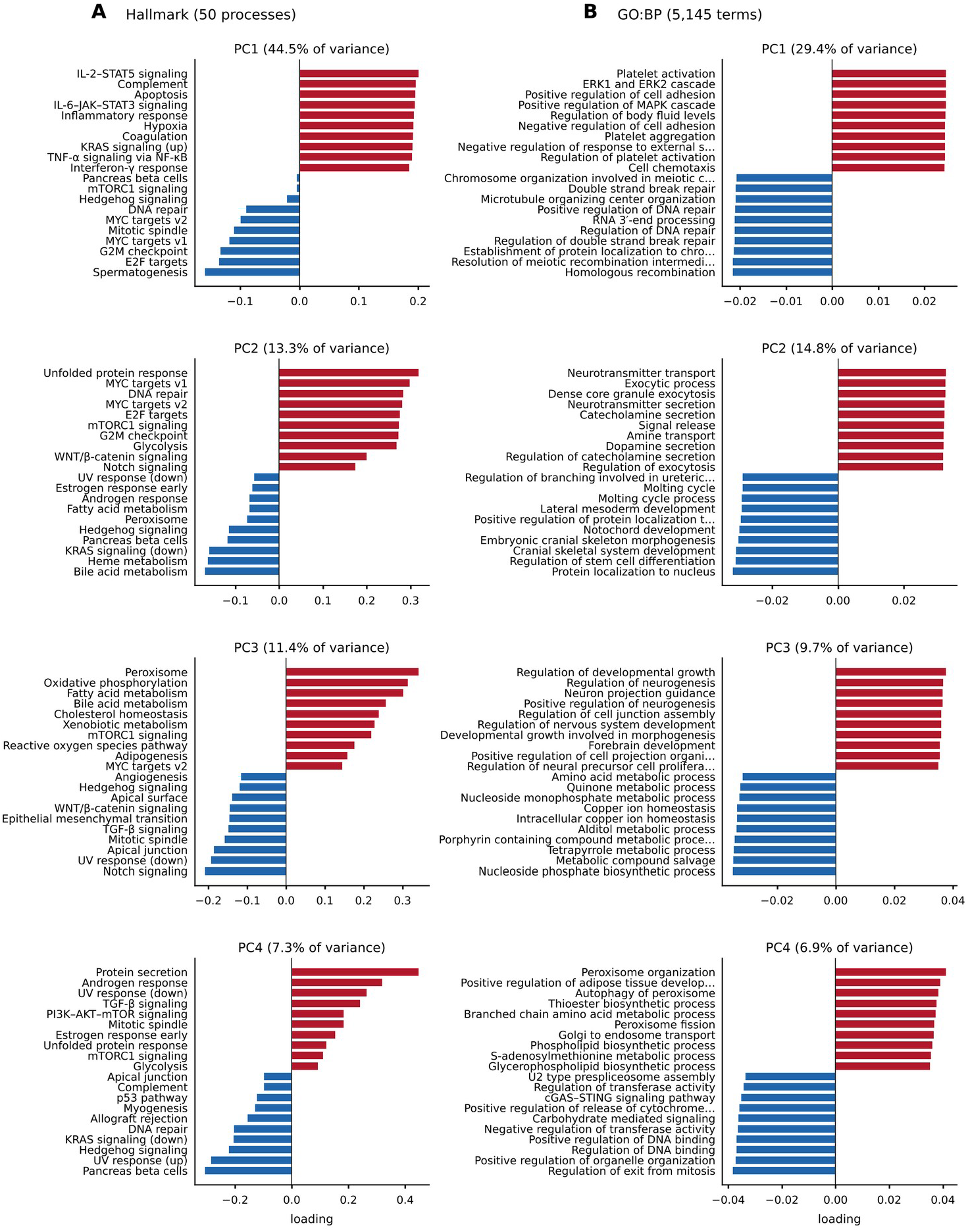
Loadings of the leading components. The ten largest loadings at each end of PC1–PC4 in the hallmark (A) and GO:BP (B) decompositions of all 901 samples, with the variance explained by each component. Red, positive end; blue, negative end. Values are in Tables S3 and S14.

**Supplementary Fig. S3.**
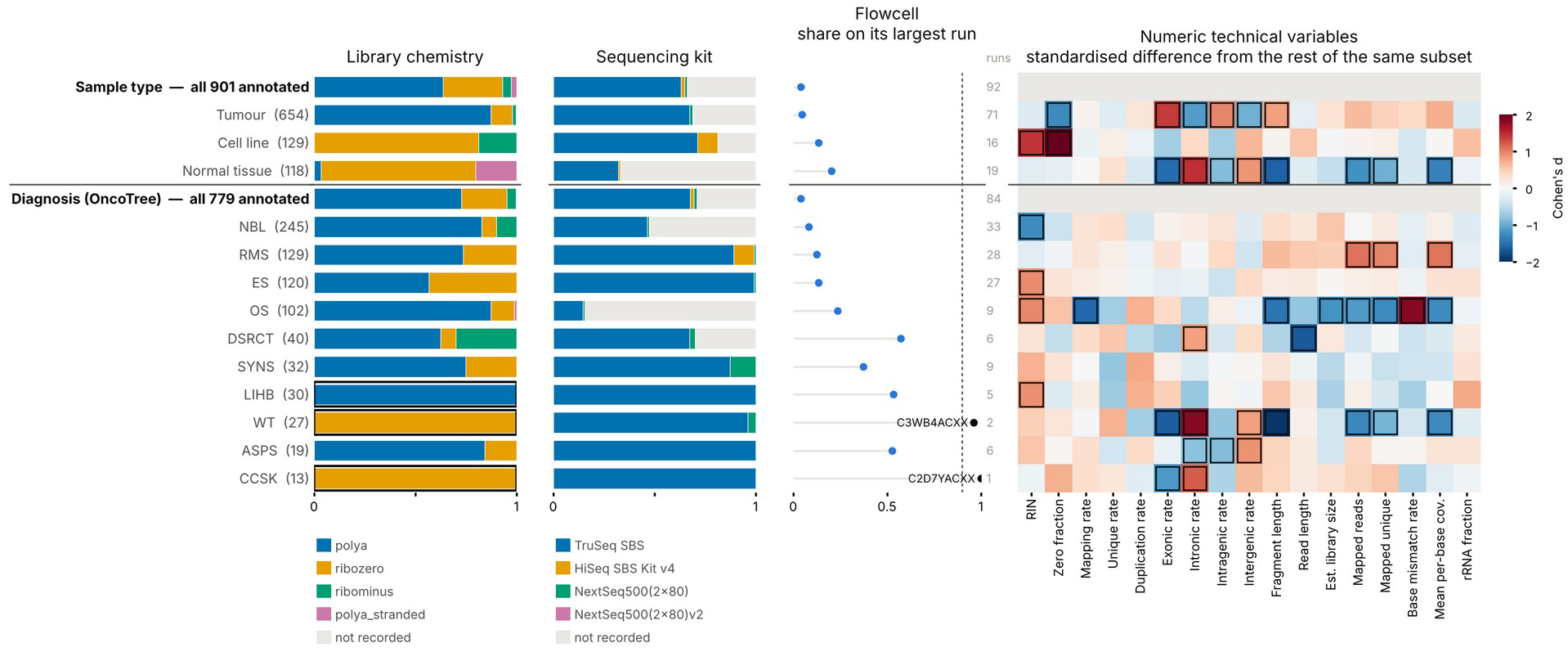
Distribution of sample types and diagnoses across technical variables. Each level with at least eight samples (rows) is shown against the samples in which its variable is recorded (bold rows). Left, composition by library chemistry and sequencing kit; centre, share of the level on its most frequent flowcell, with the number of flowcells; right, Cohen’s d of each continuous technical variable against the rest of the subset. Outlined cells mark at least 90% of a level in one stratum where the subset is below 90%, or |d| ≥ 0.8. Wilms tumour (WT) and clear cell sarcoma of the kidney (CCSK) were sequenced only with ribosomal RNA depletion, and hepatoblastoma (LIHB) only with poly(a) selection. Diagnosis codes as in Supplementary Fig. S1. Values are in Table S5.

**Supplementary Fig. S4.**
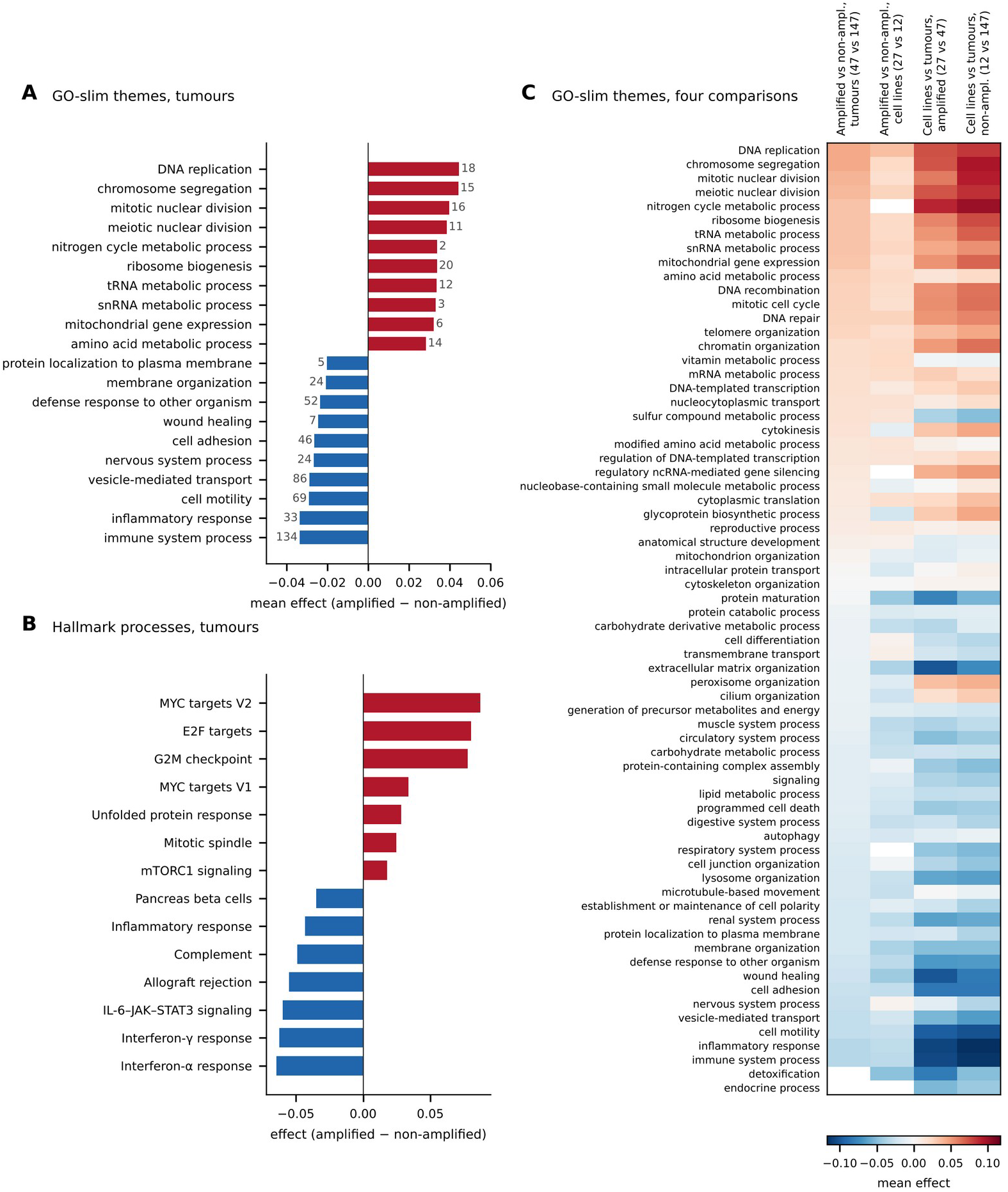
Differential pathway enrichment by *MYCN* amplification in neuroblastoma. (A) The ten GO-slim themes with the largest mean effect in each direction for amplified versus non-amplified tumours (47 versus 147); numbers give significant member terms. (B) The seven hallmark processes with the largest effects in each direction (FDR < 0.05) for the same comparison. (C) Mean effect of significant member terms for each GO-slim theme in four comparisons: amplified versus non-amplified tumours and cell lines, and cell lines versus tumours among amplified and non-amplified samples; white cells, no significant terms. Rows are ordered by the first comparison. Values are in Tables S9–S11.

**Supplementary Fig. S5.**
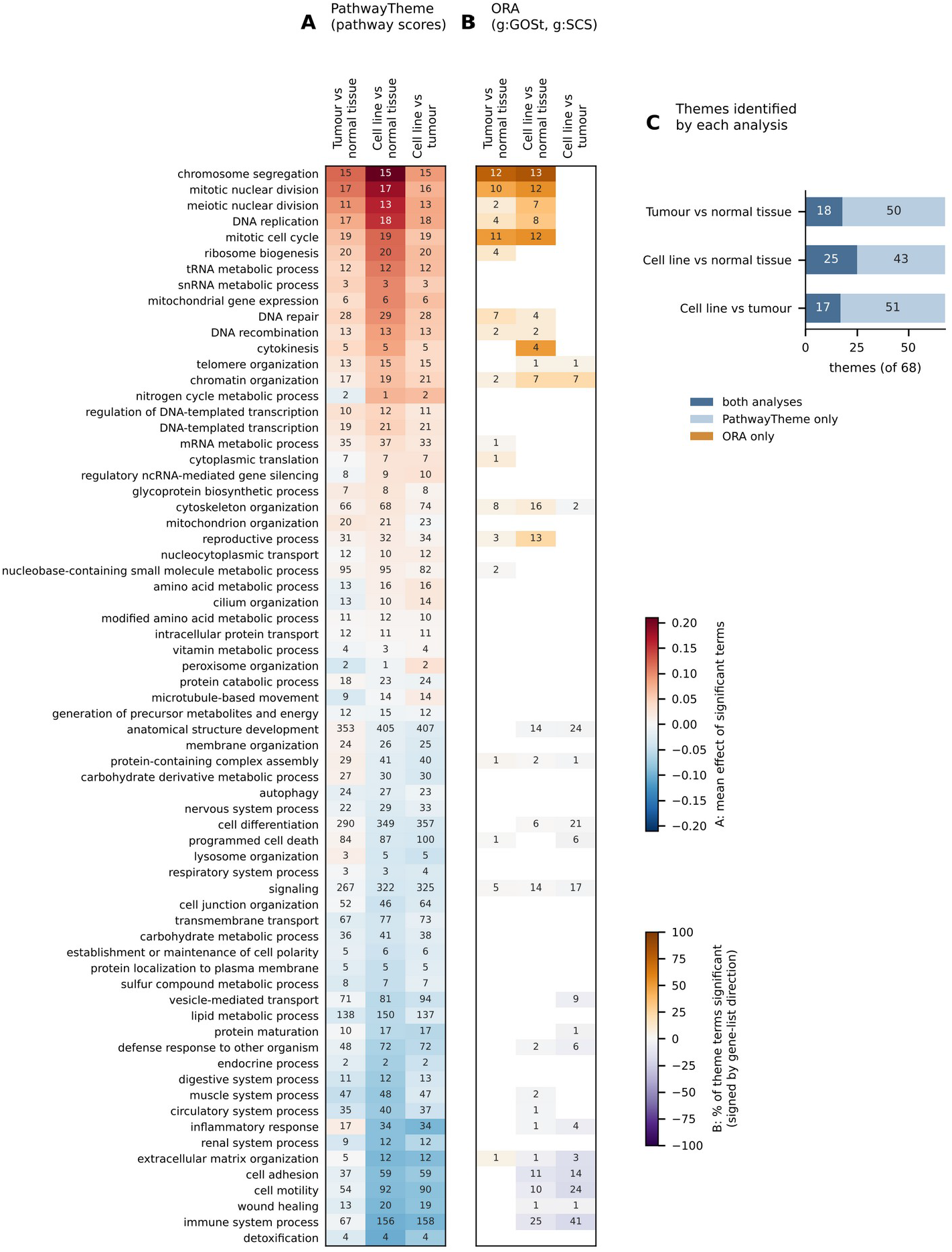
Comparison of PathwayTheme with over-representation analysis (ORA) of thresholded gene lists for the three pairwise sample-type contrasts. (A) PathwayTheme: mean effect of the significant GO:BP terms in each of the 68 GO-slim themes; numbers give significant terms. (B) ORA with g:GOSt and g:SCS correction on genes with FDR < 0.05 and | log_2_ fold change| > 2, submitted separately by direction: percentage of each theme’s terms called significant, signed by the direction of the gene list; numbers give significant terms. White cells, no significant terms. Rows are in the same order in both panels, by mean PathwayTheme effect across contrasts. (C) Number of themes identified by both analyses, by PathwayTheme only and by ORA only (none). Values are in Table S12.

## Supplementary Tables

Tables S1–S17 are provided as sheets of the accompanying workbook (PathwayTheme_Supplementary_Tables.xlsx).

**Supplementary Table S1.** Sample annotation for the 901 analysed transcriptomes. One row per sample: run-sheet fields (diagnosis, sample type, library type, RIN, sequencing kit, flowcell and identifiers), the fraction of protein-coding genes at zero TPM, the source of each completed field, the diagnosis, diagnosis code and tissue type from Supplementary Table S1 of Brohl et al., and the MYCN status read from that diagnosis code. contrast_group defines the matched tumour and cell-line groups of Fig. 1E.

**Supplementary Table S2.** Association of GO:BP components PC1–PC10 with six sample variables (Fig. 1B). For each of the 60 tests: test, number of samples and levels, statistic, effect size (rank-based η^2^ for categorical and ρ^2^ for continuous variables), P-value, Benjamini–Hochberg FDR across all 60 tests, and the level with the highest mean component score.

**Supplementary Table S3.** Loadings of the GO:BP decomposition of all 901 samples. Two selections are given: the ten largest loadings at each end of PC1–PC4 (plotted in Supplementary Fig. S2), and the largest loadings at each end of PC1–PC10 as reported by the package. Loadings are elements of unit-length component vectors; the sign of a component is arbitrary.

**Supplementary Table S4.** Association of sample type with PC1 and PC4 within each library chemistry, in both collections. Rows give the association of each component with library type (four recorded chemistries), with chemistry collapsed to two strata (poly(A) selection and ribosomal RNA depletion), and with sample type in all samples and within each stratum. The effect size in this table is the uncorrected η^2^ (between-group sum of squares over total sum of squares of the component scores), not the rank-based η^2^ of Table S2 and Fig. 1B; unconditional values therefore differ slightly from Tables S2 and S13. P-values are from the Kruskal–Wallis test, with Benjamini–Hochberg FDR across the 20 rows.

**Supplementary Table S5.** Level-wise confounding of sample type and diagnosis with 20 technical variables (Supplementary Fig. S3). Each level with at least eight samples is compared with the samples in which its own variable is recorded (reference rows, is_reference_row = True). For categorical technical variables, the value is the share of the level in its largest stratum; for continuous variables, Cohen’s d against the rest of the subset. confounded = True marks a share of at least 0.90 where the subset baseline is below 0.90, or |d| ≥ 0.8.

**Supplementary Table S6.** Separation of each diagnosis by each GO:BP component (Fig. 1C). For each diagnosis: AUC on PC1–PC10 from the one-versus-rest Mann–Whitney test among tumours and cell lines, number of samples, best-separating component, its AUC and FDR, separation as max(AUC, 1 − AUC), and the corresponding separation in the hallmark decomposition.

**Supplementary Table S7.** One-versus-rest tests of every level of each categorical variable against every GO:BP component. Mann–Whitney test, with the rank-biserial correlation as effect size and its rescaling to [0, 1] as AUC; mean component scores of the level and of the remaining samples; Benjamini–Hochberg FDR; and the end of the component occupied by the level.

**Supplementary Table S8.** GO-slim theme summaries for all GO:BP contrasts (Fig. 1E). For each analysis, contrast and theme, separately for significant (FDR < 0.05) and non-significant member terms: mean effect, number of terms, and numbers of terms increased and decreased. Effects are differences in mean ssGSEA score, case minus reference.

**Supplementary Table S9.** GO-slim themes for *MYCN*-amplified versus non-amplified neuroblastoma (Supplementary Fig. S4A). Mean effect of significant member terms in tumours (47 versus 147) and in cell lines (27 versus 12), with the rank of each theme in each comparison and the number of significant terms in tumours; immune marks themes classed as immune-related.

**Supplementary Table S10.** Differential hallmark scores for *MYCN*-amplified versus non-amplified neuroblastoma tumours (47 versus 147; Supplementary Fig. S4B). Moderated t-test; group means, effect, statistic, P-value, Benjamini–Hochberg FDR and direction for each of the 50 hallmark processes.

**Supplementary Table S11.** GO-slim theme effects for the four comparisons in the neuroblastoma cohort (Supplementary Fig. S4C): amplified versus non-amplified within tumours and within cell lines, and cell lines versus tumours within amplified and within non-amplified samples.

**Supplementary Table S12.** Comparison with over-representation analysis for the three sample-type contrasts (Supplementary Fig. S5). For each contrast and theme: mean effect and number of significant member terms from PathwayTheme; and the number of member terms called by g:GOSt with g:SCS correction, with the percentage of the theme’s terms called, signed by the direction of the gene list in which they were called.

**Supplementary Table S13.** Association of hallmark components PC1–PC10 with six sample variables; as Table S2, for the hallmark collection (50 processes).

**Supplementary Table S14.** Loadings of the hallmark decomposition of all 901 samples; as Table S3.

**Supplementary Table S15.** One-versus-rest tests of every level of each categorical variable against every hallmark component; as Table S7.

**Supplementary Table S16.** Agreement of the moderated t-test in PathwayTheme with limma::eBayes (Supplementary Fig. S1). For each of 24 contrasts from four analyses: group sizes, number of pathways, prior degrees of freedom and prior variance from both implementations with their relative difference, maximum absolute differences in effect, moderated t and P-value, Pearson correlation of the moderated t values, and the number of pathways whose significance call differs.

**Supplementary Table S17.** All contrasts tested, with group sizes and the numbers of pathways tested, significant (FDR < 0.05), increased and decreased. Contrasts include diagnoses with fewer than ten samples, which are not interpreted in the main text.

## References

1. Leek JT, Scharpf RB, Bravo HC et al. Tackling the widespread and critical impact of batch effects in high-throughput data. Nat Rev Genet 2010;11:733–739.

2. Hicks SC, Townes FW, Teng M et al. Missing data and technical variability in single-cell RNA-sequencing experiments. Biostatistics 2018;19:562–578.

3. Blighe K, Andrews J, Lun A. PCAtools: everything principal components analysis. R/Bioconductor package. doi:10.18129/B9.bioc.PCAtools.

4. McCarthy DJ, Campbell KR, Lun ATL et al. Scater: pre-processing, quality control, normalization and visualization of single-cell RNA-seq data in R. Bioinformatics 2017;33:1179–1186.

5. Li J, Bushel PR, Chu T-M et al. Principal variance components analysis: estimating batch effects in microarray gene expression data. In: Scherer A (ed.), Batch Effects and Noise in Microarray Experiments: Sources and Solutions. Chichester: Wiley, 2009, 141–154.

6. Reese SE, Archer KJ, Therneau TM et al. A new statistic for identifying batch effects in high-throughput genomic data that uses guided principal component analysis. Bioinformatics 2013;29:2877–2883.

7. Marini F, Binder H. pcaExplorer: an R/Bioconductor package for interacting with RNA-seq principal components. BMC Bioinformatics 2019;20:331.

8. Brohl AS, Sindiri S, Wei JS et al. Immuno-transcriptomic profiling of extracranial pediatric solid malignancies. Cell Rep 2021;37:110047.

9. Frost HR. Variance-adjusted Mahalanobis (VAM): a fast and accurate method for cell-specific gene set scoring. Nucleic Acids Res 2020;48:e94.

10. Holland CH, Tanevski J, Perales-Patón J et al. Robustness and applicability of transcription factor and pathway analysis tools on single-cell RNA-seq data. Genome Biol 2020;21:36.

11. Wang RH, Thakar J. Comparative analysis of single-cell pathway scoring methods and a novel approach. NAR Genom Bioinform 2024;6:lqae124.

12. Barbie DA, Tamayo P, Boehm JS et al. Systematic RNA interference reveals that oncogenic KRAS-driven cancers require TBK1. Nature 2009;462:108–112.

13. Hänzelmann S, Castelo R, Guinney J. GSVA: gene set variation analysis for microarray and RNA-seq data. BMC Bioinformatics 2013;14:7.

14. Foroutan M, Bhuva DD, Lyu R et al. Single sample scoring of molecular phenotypes. BMC Bioinformatics 2018;19:404.

15. Wieder C, Lai RPJ, Ebbels TMD. Single sample pathway analysis in metabolomics: performance evaluation and application. BMC Bioinformatics 2022;23:481.

16. Ge SX, Son EW, Yao R. iDEP: an integrated web application for differential expression and pathway analysis of RNA-Seq data. BMC Bioinformatics 2018;19:534.

17. Frost HR, Li Z, Moore JH. Principal component gene set enrichment (PCGSE). BioData Min 2015;8:25.

18. Wagner F. GO-PCA: an unsupervised method to explore gene expression data using prior knowledge. PLoS One 2015;10:e0143196.

19. DeTomaso D, Jones MG, Subramaniam M et al. Functional interpretation of single cell similarity maps. Nat Commun 2019;10:4376.

20. Odom GJ, Ban Y, Colaprico A et al. PathwayPCA: an R/Bioconductor package for pathway based integrative analysis of multi-omics data. Proteomics 2020;20:e1900409.

21. Tomfohr J, Lu J, Kepler TB. Pathway level analysis of gene expression using singular value decomposition. BMC Bioinformatics 2005;6:225.

22. Fan J, Salathia N, Liu R et al. Characterizing transcriptional heterogeneity through pathway and gene set overdispersion analysis. Nat Methods 2016;13:241–244.

23. Meng C, Basunia A, Peters B et al. MOGSA: integrative single sample gene-set analysis of multiple omics data. Mol Cell Proteomics 2019;18:S153–S168.

24. Supek F, Bošnjak M, Škunca N et al. REVIGO summarizes and visualizes long lists of Gene Ontology terms. PLoS One 2011;6:e21800.

25. Sayols S. rrvgo: a Bioconductor package for interpreting lists of Gene Ontology terms. microPublication Biology 2023. doi:10.17912/micropub.biology.000811.

26. Gu Z, Hübschmann D. simplifyEnrichment: a Bioconductor package for clustering and visualizing functional enrichment results. Genomics Proteomics Bioinformatics 2023;21:190–202.

27. Fang Z, Liu X, Peltz G. GSEApy: a comprehensive package for performing gene set enrichment analysis in Python. Bioinformatics 2023;39:btac757.

28. Chen EY, Tan CM, Kou Y et al. Enrichr: interactive and collaborative HTML5 gene list enrichment analysis tool. BMC Bioinformatics 2013;14:128.

29. Klopfenstein DV, Zhang L, Pedersen BS et al. GOATOOLS: a Python library for Gene Ontology analyses. Sci Rep 2018;8:10872.

30. Pedregosa F, Varoquaux G, Gramfort A et al. Scikit-learn: machine learning in Python. J Mach Learn Res 2011;12:2825–2830.

31. Benjamini Y, Hochberg Y. Controlling the false discovery rate: a practical and powerful approach to multiple testing. J R Stat Soc Series B 1995;57:289–300.

32. Kerby DS. The simple difference formula: an approach to teaching nonparametric correlation. Compr Psychol 2014;3:11.IT.3.1.

33. Smyth GK. Linear models and empirical Bayes methods for assessing differential expression in microarray experiments. Stat Appl Genet Mol Biol 2004;3:Article 3.

34. Ritchie ME, Phipson B, Wu D et al. limma powers differential expression analyses for RNA-sequencing and microarray studies. Nucleic Acids Res 2015;43:e47.

35. Ashburner M, Ball CA, Blake JA et al. Gene Ontology: tool for the unification of biology. Nat Genet 2000;25:25–29; Gene Ontology Consortium. The Gene Ontology knowledgebase in 2023. Genetics 2023;224:iyad031.

36. Menon SV, Gom A, Gillmor AH et al. Longitudinal profiling of ASPSCR1::TFE3 Type 2 alveolar soft part sarcoma unveils therapeutic targets. EMBO Mol Med 2026 (in press).

37. Liberzon A, Birger C, Thorvaldsdóttir H et al. The Molecular Signatures Database hallmark gene set collection. Cell Syst 2015;1:417–425.

38. Kolberg L, Raudvere U, Kuzmin I et al. g:Profiler—interoperable web service for functional enrichment analysis and gene identifier mapping (2023 update). Nucleic Acids Res 2023;51:W207–W212.

## Supplementary references

S1. Frankish A, Diekhans M, Jungreis I et al. GENCODE 2021. Nucleic Acids Res 2021;49:D916–D923.

S2. Liberzon A, Birger C, Thorvaldsdóttir H et al. The Molecular Signatures Database hallmark gene set collection. Cell Syst 2015;1:417–425.

S3. Ashburner M, Ball CA, Blake JA et al. Gene Ontology: tool for the unification of biology. Nat Genet 2000;25:25–29.

S4. Cohen J. Statistical Power Analysis for the Behavioral Sciences, 2nd edn. Hillsdale, NJ: Lawrence Erlbaum, 1988. Numbered references [n] refer to the main-text reference list.

